# High-resolution mass spectrometry-based non-targeted metabolomic discovery of disease and glucocorticoid biomarkers in an animal model of muscular dystrophy

**DOI:** 10.1101/475590

**Authors:** Mathula Thangarajh, Simina M. Boca, Aiping Zhang, Kirandeep Gill, Habtom Ressom, Zhenzhi Li, Eric P. Hoffman, Kanneboyina Nagaraju, Yetrib Hathout

## Abstract

Urine is increasingly being considered as a source of biomarker development in Duchenne Muscular Dystrophy (DMD), a severe, life-limiting disorder that affects approximately 1 in 4500 boys. In this study, we used the mdx mice—a murine model of DMD—to discover biomarkers of disease, as well as pharmacodynamic biomarkers responsive to prednisolone, commonly used to treat DMD. Longitudinal urine samples were analyzed from male age-matched mdx and wild-type mice randomized to prednisolone or vehicle. We used high-resolution mass spectrometry to discover metabolic biomarkers of both disease and glucocorticoid treatment. A large number of metabolites (869 out of 6,334) were found to be significantly different between mdx and wild-type mice at baseline (Bonferroni-adjusted p-value < 0.05), thus being associated with disease status. These included a peak with m/z=357 and creatine, which were also discovered in a previous human study looking at serum. Novel observations included biliverdin and hypusine. These four peaks were also significantly higher in mdx mice compared to wild-type, as well as significantly associated with time after the baseline. Creatine and biliverdin were also associated with treatment after the baseline, but the association with creatine may have been driven by an imbalance at baseline. In conclusion, our study reports a number of biomarkers, both known and novel, which may be related to either the mechanisms of muscle injury in DMD and/or prednisolone treatment.

## Introduction

Oral corticosteroids are the standard of care in Duchenne Muscular Dystrophy (DMD)—an X-linked genetic disease that affects 1 in 4000 to 5000 boys worldwide [1,2]. DMD is caused by mutations in the dystrophin gene located on the short arm of the × chromosome [3,4]. Oral corticosteroid use in young boys with DMD changes the natural history of the disease. One of the largest natural history studies in DMD showed that corticosteroids increased survival in DMD, improved motor outcomes such as continued ambulation, as well as cardiopulmonary outcomes [5]. Despite these unambiguous clinical benefits, there is great variation in the dosage and regimen of corticosteroids, and with compliance to treatment in DMD [6]. Patients and families often discontinue corticosteroids mainly due to unpleasant side-effects including sleep disturbance, weight gain, increased fracture risk, and metabolic syndrome. Identification of non-invasive pharmacodynamic biomarkers associated with glucocorticoid side effects in boys may help predict those individuals who might be at a higher risk for developing corticosteroid-induced side-effects [7].

We have previously identified a metabolic signature consisting of serum biomarkers in DMD [8]. Metabolites that were increased in individuals with DMD compared to healthy controls included creatine, arginine, and unknown compounds with m/z values of approximately 357 and 312 Da; metabolites that were decreased in DMD included creatinine, androgen derivatives, and other unknown, yet-to-be-identified metabolites. In order to better characterize these metabolites as potential biomarkers in DMD, we used the *mdx* mouse—a widely used pre-clinical model—to agnostically evaluate urine metabolites associated with dystrophin deficiency and/or response to corticosteroids. Urine is an easily accessible bodily fluid, can be collected non-invasively and on multiple time-points, and is becoming an attractive source of biomarker development in DMD [9–12]. The identification of a palette of pharmacodynamic urine metabolites will allow the identification of metabolic pathways that are affected by the disease pathogenesis and that might be responsive to therapies such as corticosteroids and exon skipping. Such pharmacodynamic biomarkers may also help in drug development and dose finding.

## Methods

### Animals

The experiments using mice described in this paper were conducted according to the institutional guidelines regarding the humane treatment of animals. This study was done in compliance with the National Institutes of Health guidelines for pre-clinical studies as described by [13] and the standard operating procedure for pre-clinical studies [14]. The protocol was approved by the local Institutional Animal Care and Use Committee (#30424).

Age-matched 5-week-old male wild-type (C57BL/10ScSnJ) and *mdx* (C57BL/10ScSn-*Dmd*<*mdx*>/J) mice litters were purchased from Jackson Laboratory (Bar Harbor, ME). Mice were acclimatized for 4 days prior to commencement of experiments. A total of 48 mice (23 wild-type (WT) and 25 *mdx*) were used in order to ensure a rigorous statistical analysis. WT and *mdx* mice matched for body weight were randomized into treatment with either vehicle control (cherry syrup) or prednisolone (dissolved in cherry syrup) for a total of 4 weeks as previously published [15]. Animals were treated with vehicle or prednisolone (5mg/kg/day) by mouth based on body weight. Animals had access to water and food ad libitum. The study team was blinded to the treatment groups. Unblinding of the different treatment groups occurred after data collection and data processing.

### Urine collection

Urine from mice was collected by placing them in clean cages early in the morning. Urine was collected at three different time points: baseline (prior to treatment), 15 days following commencement of treatment, and at completion of treatment at 4 weeks; these are coded as times T0, T1, and T2 in the remainder of this manuscript. We note that at T0, the mice had not received any vehicle cherry syrup. The urine collection had the following caveats: 4 *mdx* and 2 WT mice died between times T0 and T1, and some mice did not urinate at every time point; one sample was not available for T2 and therefore not analyzed. In total, 48 mice were evaluated at time T0 and 43 unique mice at times T1 and T2, out of which 41 provided measurements at T1 but not T2 and 41 provided measurements at T2 but not T1.

### LC-MS method

For metabolite extraction, 80 µl of a solution of 50% acetonitrile in water containing internal standards (10µl of 1mg/ml debrisoquine and 50µl of 1mg/ml 4-nitrobenzoic acid added to 10 ml of 50% acetonitrile in water) was added to 20 µl of each urine sample in an Eppendorf vial. The samples were centrifuged at 13,000 rpm for 20 minutes at 4°C and the supernatant transferred to fresh vials for UPLC-Qtof analysis. 2µl of each sample was injected onto a Waters Acquity BEH C18 1.7 µm, 2.1 × 50 mm column using an Acquity UPLC system by Waters Corporation, Milford, MA. The gradient mobile phase consisted of solvent A - 100% water with 0.1% formic acid - and solvent B - 100% acetonitrile with 0.1% formic acid. The column temperature was set to 40° C and flow rate to 0.5 ml/min. The gradient started with 95% of solvent A and shifted to 80% of solvent A at 4 minutes at a ramp of curve 6. From 4.0 to 8.0 minutes, the gradient moved to 95% of solvent B and stayed there until 9 minutes. It returned to initial conditions of 95% solvent A and 5% solvent B at 11 minutes. The elution from the column was introduced to a quadrupole time of flight mass spectrometer (Waters G2-Qtof) by electrospray ionization in both positive and negative mode at a capillary voltage of 3.0 kV and sampling cone voltage of 30 V. The source temperature was set to 120° C and the desolvation temperature to 500 °C. The cone gas flow was maintained at 25L/hr and desolvation gas flow at 1000L/hr. Leucine-encephalin solution in 50% acetonitrile was used a reference mass ([M+H]^+^ = 556.2771 and [M-H]^−^ = 554.2615). The data were acquired in centroid mode from mass range of 50 to 1200 with the software Mass lynx (Waters Corporation). Pooled quality controls samples were injected after every 10 injections.

### Data processing

LC-MS metabolomics samples were preprocessed as previously described [8]. In brief, the XCMS approach [16], available through the xcms package on Bioconductor [17] was used to detect features and estimate intensities, perform retention time correction, and group peaks from different samples, followed by filling in missing peaks via integrating the signal in the peak region defined by the other samples. The resulting peaks were then annotated using the CAMERA package [18], resulting in the prediction of possible isotopes.

Following these initial preprocessing steps, peak groups with fewer than 37 peaks were removed, along with predicted isotopes. This led to 3,674 peaks in the positive mode and 2,663 peaks in the negative mode. Internal standard normalization was then performed, with the intensities in the each mode being divided by the corresponding standard and the standards being removed from the list of peaks, leading to a total number of 6,335 peaks. A small number of peak-sample combinations had intensities of 0 (51 for the negative mode and 3 for the positive mode); these intensities were replaced by the smallest non-zero intensity for that peak across all samples. The intensities were then log2-transformed and the peaks from both modes were quantile-normalized together [19]. All the quality control samples were removed from the downstream analysis.

### Statistical analyses

Given that the mice had a different diet at time T0 compared to times T1 and T2, the analyses considered were: 1) a comparison of the genotypes at time T0 via a two-sample t-test and 2) a mixed effects linear model for times T1-T2 which included genotype, time, and treatment, as well as all their interactions, with a random intercept to account for the within-mouse correlation. We note that for analysis 1) we did not consider treatment given that all the mice were at baseline, with none being yet treated with prednisolone. Prior to analysis 1), a single peak that had the same value in all samples at T0 was removed, so that this analysis was performed on 6,334 peaks. The correlation between the p-values for t-tests assuming equal variances and t-tests assuming difference variances in the two groups was > 0.9999; moving forward, we used the results from the analysis with the equal variance assumption. For analysis 2), we focused on the top 50 most significant peaks from analysis 1), with the addition of creatine and creatinine, which are well-known to be important in DMD and were among the top metabolites associated with disease status in previous work [8]. In the mixed effects models, likelihood ratio tests were used to assess the association of genotype, time, and treatment with the transformed peak intensities. In order to account for multiple testing, statistical significance was determined to be reached if the Bonferroni-adjusted p-values were less than 0.05, when considering all 6,334 peaks.

### Validation of top peaks by LC-MS/MS

The top 50 peaks in terms of significant differences between *mdx* and WT mice at baseline were considered as queries in the Human Metabolome Database (HMDB), looking for possible H+, Na+, and K+ adducts for the positive mode peaks and H-adducts for the negative mode peaks, within 10 ppm accuracy. Eleven of these peaks were selected to be validated by MS/MS. Most of them were also significantly different between the genotypes longitudinally. Creatine and creatinine were previously validated, so we did not include them for validation by the MS/MS method here. The positive mode peaks considered in the MS/MS experiment had the following m/z values: 229.16, 234.18, 357.25, 485.33, 583.26, 705.18, 884.93 Da. The negative mode peaks considered in the MS/MS experiment had the following m/z values: 210.11, 486.16, and 652.11 Da. The validation experiments were similar to those in [8], running the samples on Waters G2-Qtof. Pooled QC samples were processed in a similar manner to the profiling experiment and the chromatographic conditions used for data acquisition remained the same. The MS/MS spectra obtained were then inspected by matching to the online available spectra from the databases Metlin [20] and HMDB [21] and manually. Further putative identification was performed as follows: The MZXML files were parsed using the *pyteomics* python library [22]. For each sample, the retention time was used to access the scan of the targeted compound. The mzxml function in *pyteomics* was used to access the m/z and corresponding intensity information of the retention time. The top 30 peaks with highest intensities were used for compound identification using MS/MS spectral libraries such as Metlin, HMDB, and MONA (http://mona.fiehnlab.ucdavis.edu/). In addition to searching tools provided by these sources, we used MetaboSearch Pro [23] to search against these libraries and assign scores to the putative metabolite IDs.

## Results

### Comparison of genotypes at baseline

Out of a total of 6,334 detected peaks, 868 were determined to show significant differences between the *mdx* and the WT mice (Bonferroni-adjusted p-value < 0.05). Given this large number, for the downstream analyses, we chose to focus on the top 50 most significant peaks, along with creatine (m/z = 132.08) and creatinine (m/z = 114.06), as described above. A subset of these 50 peaks was also selected for validation via MS/MS; the MS/MS spectra for the 11 selected peaks considered in this follow-up experiment are in Supplemental Figure 1. Creatine and creatinine are known to be important in DMD and were among the top peaks in our human serum study [8]. We note that the positive mode peak, with m/z = 357.25 (p = 5.11 × 10^−25^ unadjusted), appears to be the same as the top peak in our previous work with DMD patients [8]. Interestingly, creatine was not among the top 50 peaks in the mouse model, but ranked 472 and was still deemed significant (p = 4.91 × 10^−8^ unadjusted). Creatinine, however, was not significant (p = 0.98 unadjusted). Boxplots representing results for these 3 peaks for the comparison between genotype are shown in Figure 1a-c. We considered two additional peaks at m/z = 234.18 and m/z = 583.26 which were significant (p = 1.26 × 10^−14^ unadjusted and p = 2.31 × 10^−16^ unadjusted, respectively) and validated using MS/MS in Figure 1 d-e. The peak at m/z = 234.18 was identified as hypusine and the peak at m/z = 583.26 was identified as a geometric isomer of biliverdin in accordance with recently published MS/MS data of biliverdin [24]. Peaks putatively identified using the combined HMDB+Metlin+MONA+MetaboSearch Pro are listed in Supplemental Table 2. For all peaks in Figure 1 besides creatinine, the intensities are significantly elevated in the urine of *mdx* mice compared to WT mice. In particular, for the peak with m/z = 357.25, this finding recapitulates our earlier findings from the serum of younger individuals with DMD. For creatine, older boys with DMD had higher values [8]. We note that the only match for this peak is a tripeptide, Pro Ile Gln; however, the MS/MS spectrum did not appear to have a perfect fragmentation pattern for a tripeptide perhaps due to overlapping contaminants. Upon matching this tripeptide against the *Mus musculus* proteome using only UniProt/SwissProt and not including isoforms at https://research.bioinformatics.udel.edu/peptidematch/index.jsp [25], we obtained 770 proteins, including the mouse titin protein, in which it appears 7 times. Titin fragments have been previously detected in urine samples collected from DMD patients [10,26–28].

**Figure 1.**
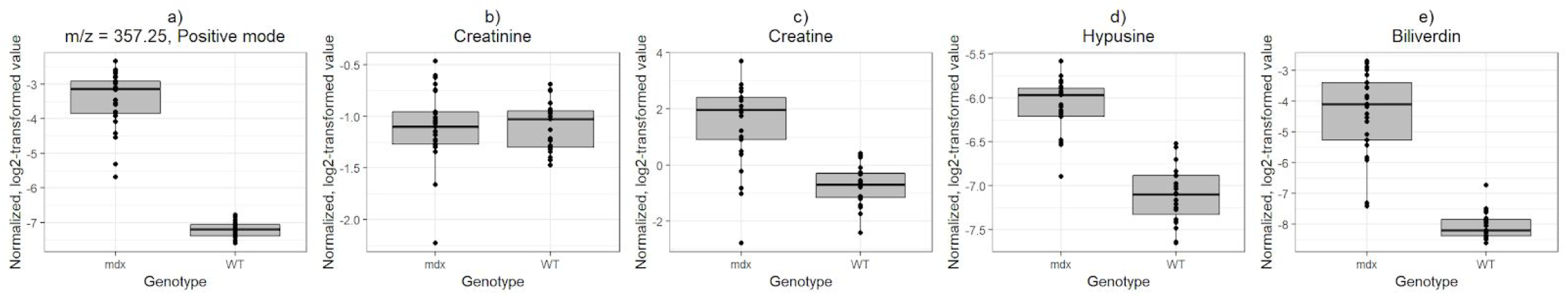
Genotype comparison of normalized, log2-transformed intensity values at baseline (T0) for a) peak with m/z = 357.25, b) creatine (m/z = 132.08 in positive mode), c) creatinine (m/z = 114.06 in positive mode), d) hypusine (m/z = 234.18 in positive mode), e) biliverdin (m/z = 583.26 in positive mode).

### Comparison of genotypes, time, and treatment for T1-T2

As discussed above, we focused our analysis at the later time points on the peaks found to have the lowest 50 p-values in the genotype comparison at T0, in addition to creatine and creatinine. This is because we are of the opinion that it is important to focus on peaks that are likely to be indicative of the disease process, with differences seen at an early age. Creatine and all the peaks selected for validation except for the one at m/z = 370.04 were among those associated with genotype at T1-T2. Creatine, biliverdin, and the peaks at m/z = 210.11, 484.16, 884.93, and 485.33 were significantly associated with treatment at T1-T2. Creatine and all the peaks selected for validation except for the one at m/z = 484.16 were among those associated with time. Overall, 31 of these 52 peaks were significantly associated with genotype, 6 with treatment, and 17 with time. The results from the HMDB query for the top peaks are in Supplemental Table 1. The overlaps between these categories are summarized in Figure 2.

**Figure 2.**
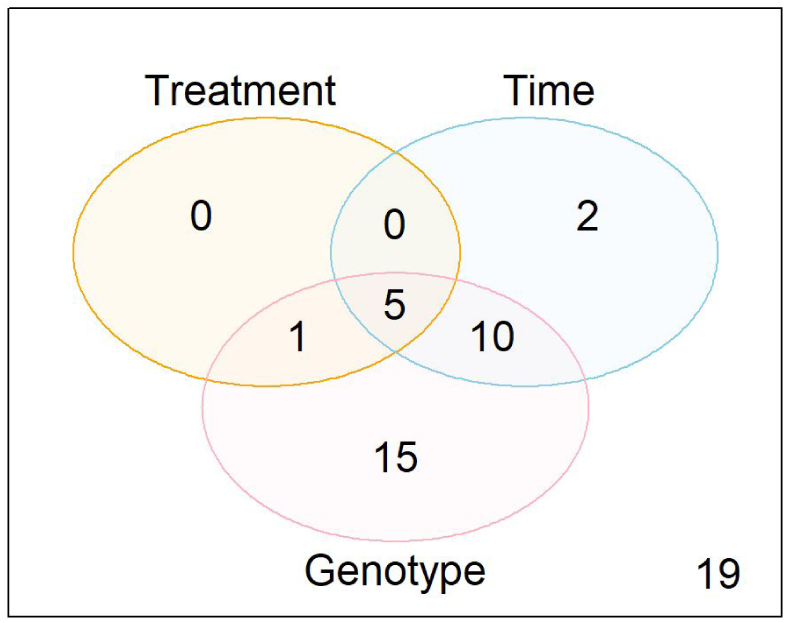
Overlaps between the peaks significantly associated with treatment, time, and genotype at times T1-T2 (Bonferroni-adjusted p-value < 0.05 for 6,334 peaks) out of the 52 peaks considered, consisting of the top 50 peaks most significantly associated with genotype at T0, plus creatine and creatinine. 19 peaks were not significantly associated with treatment, time, or genotype (bottom right corner).

In particular, we note that the peak with m/z = 357.25 was associated with genotype (p = 1.71 × 10^−40^ and time (p = 1.99 × 10^−25^ unadjusted), but not with treatment (p = 7.11 × 10^−4^ unadjusted, p = 1 when adjusting for multiple comparisons). Creatine was associated with genotype, as well as with treatment and time. However, the treatment association for creatine may be driven at least in part by an imbalance at baseline between the wild-type mice later randomized to treatment 1 versus those randomized to treatment 2 (p = 0.009). Creatinine was not significantly associated with any variable. Figure 3 shows the trends for these five metabolites. Hypusine and biliverdin were also strongly associated with genotype (p = 9.28 × 10^−9^ unadjusted, p = 1.59 × 10^−16^ respectively). Hypusine was also associated with time (p = 2.66 × 10^−8^ unadjusted), but not with treatment (p = 0.16 unadjusted). Biliverdin was associated with both treatment (p = 2.24 × 10^−6^ unadjusted) and time (p = 2.01 × 10^−14^) and did not show a major imbalance at baseline.

**Figure 3.**
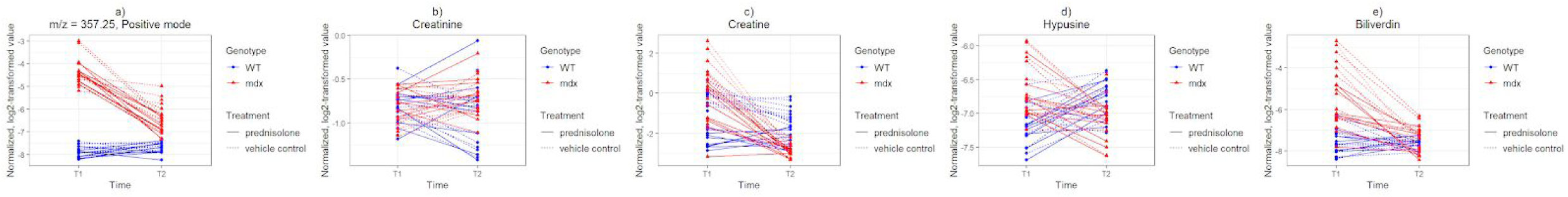
Normalized, log2-transformed intensity values at times T1-T2 for a) peak with m/z = 357.25 in positive mode, b) creatine (m/z = 132.08 in positive mode), c) creatinine (m/z = 114.06 in positive mode), d) hypusine (m/z = 234.18 in positive mode), e) peak likely to be biliverdin (m/z = 583.26 in positive mode).

Once again, the results for the peak with m/z = 357.25, showing a decrease over time in the genotype with the disorder, recapitulate the results from the human serum and urine studies [8,10,29]. An example of a peak showing treatment differences (p = 6.87 × 10^−9^) that did not have a large imbalance at baseline and that, interestingly, showed overall higher values in the *mdx* versus the WT group is the peak with m/z = 484.16, shown in Figure 4. Supplemental Figure 2 shows the baseline (T0) results for this peak.

**Figure 4.**
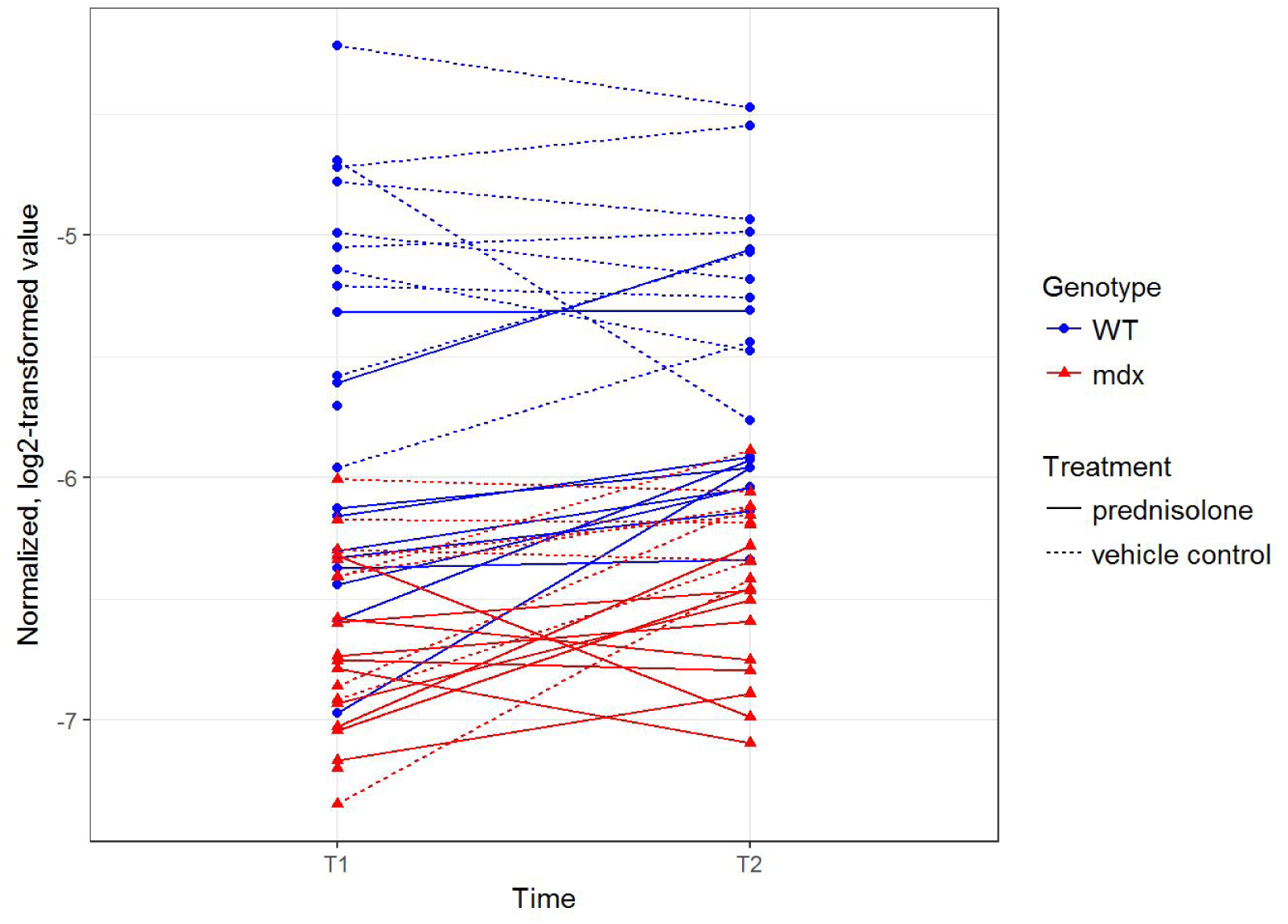
Normalized, log2-transformed intensity values at times T1-T2 for peak with m/z = 484.16 in negative mode at times T1-T2.

We also highlight a peak with m/z = 884.92, which demonstrated a significant decrease over time, as well as a significant association with treatment (p = 1.37 × 10^−8^) and a clear separation between *mdx* and WT mice in Figure 5 (boxplot for time T0 in Supplemental Figure 3).

**Figure 5.**
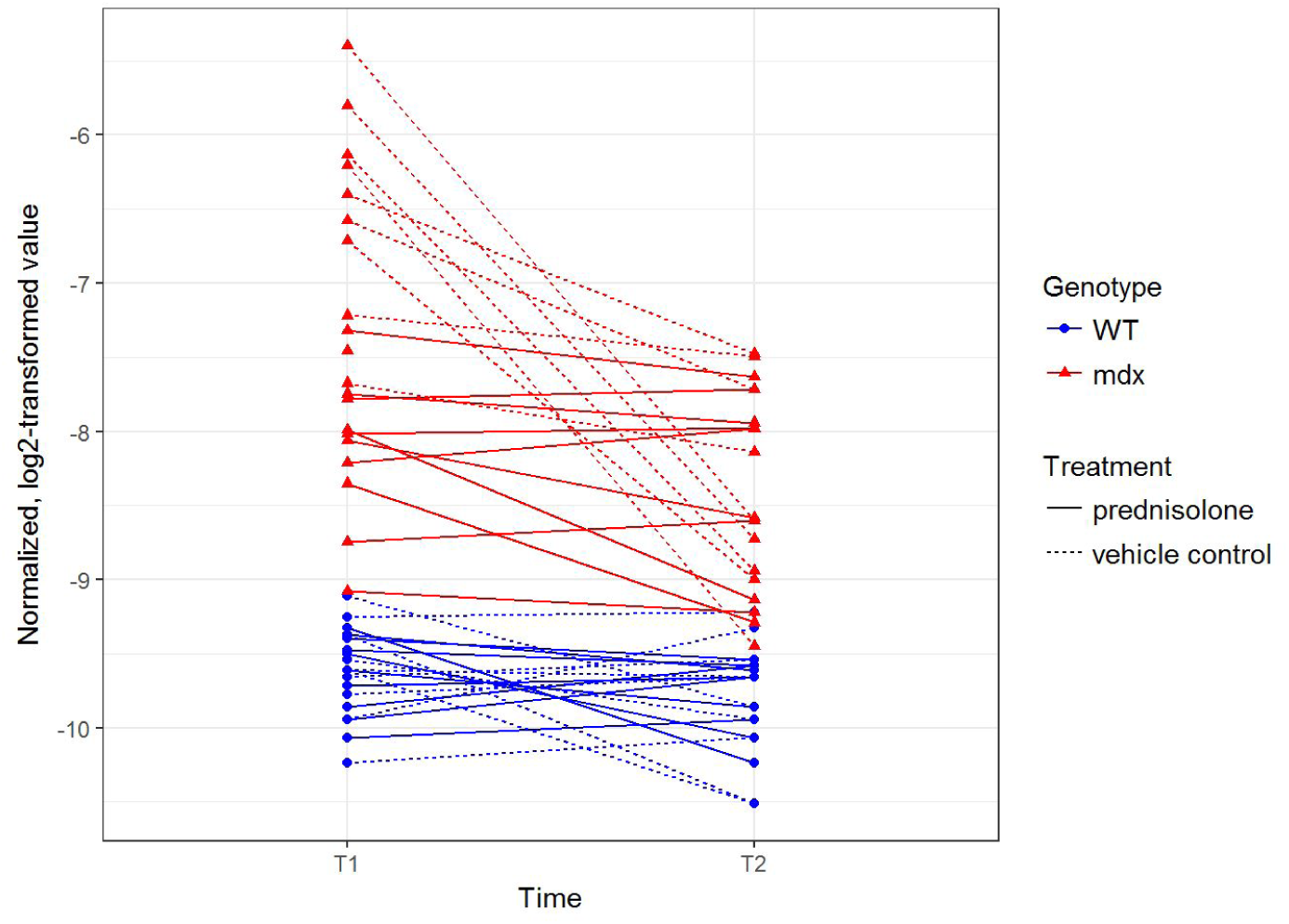
Normalized, log2-transformed intensity values at times T1-T2 for peak with m/z = 884.92 in positive mode at times T1-T2.

## Discussion

Our primary goal in this study was to identify urinary metabolite biomarkers associated with (i) muscle pathology and (ii) prednisone response in an animal model of DMD. We foresee that such information can guide future studies to evaluate biomarkers to predict disease progression and response to corticosteroids and other anti-inflammatory drugs. To our knowledge, this is the first study evaluating urine metabolites in the murine model of DMD. There is a growing body of literature that shows that urine is becoming an attractive biomarker in DMD [9–12,30], including that changes in urinary biomarker signatures may inform regarding response to therapy [30]. It is also easily accessible and less invasive, especially to very young boys.

The myriad biomarkers reported here are reflective possibly of the different mechanisms that result in muscle injury in DMD. The best-known biomarkers of DMD is creatine kinase, which seeps from muscle into blood and is generally used to identify young boys with this disorder, being many times higher than in healthy individuals [31]. While this an appropriate diagnostic biomarker, we have shown that it decreases with disease progression and muscle loss [32]. We have also previously shown that creatine and creatinine are substantially different in the serum of DMD cases compared to controls, with creatine being higher in the cases and creatinine being lower; these metabolites are both in the creatine metabolism pathway, which includes creatine kinase [8]. Our current work shows an increase in creatine at baseline in the *mdx* mice over the WT mice, although it tends to decrease over time, whereas in the human serum study, it stayed constant or slightly increased over time in DMD cases and decreased in controls. However, creatinine was not significant in any of the analyses considered. Differences in these results could be due to the biospecimen (serum vs urine), model (human disease vs murine model), or type of study (observational natural history study vs randomized laboratory study).

Particularly interesting is the metabolite corresponding to the peak with m/z = 357.25 (positive mode). This metabolite was the top metabolite in our previous study [8]. Furthermore, it recapitulates the trend of being generally higher in cases than in controls, as well as decreasing over time in cases. While we have been unable to definitively identify it herein, we consider that could be an important key to the pathogenesis of DMD and thus worthy of further investigation. If this peak is indeed the tripeptide Pro Ile Gln and it is a titin fragment, this would confirm many prior findings of elevated titin levels in both the urine and serum of young DMD patients [10,29,33], however this remains speculative at this point.

Two novel observations from our study concern biliverdin and hypusine, metabolites which were higher in *mdx* urine compared to WT mice. At times T1 and T2, both metabolites showed associations with genotype and time, with biliverdin also showing an association with treatment. Biliverdin is an end-product in the heme pathway. Heme oxygenase catabolizes heme to equimolar biliverdin, carbon monoxide, and ferrous iron. Heme oxygenase is considered a regulator of oxidative stress and is anti-inflammatory. Second, its product carbon monoxide has vasorelaxant property. Recent studies have provided contradictory information on whether heme oxygenase-1 is decreased [34,35] or increased [36,37] in the muscle of *mdx* mice compared to controls. We postulate that a pro-inflammatory state seen in *mdx* mice drives active breakdown of heme, resulting in an increase in biliverdin levels in urine [38,39]. Further investigation of this pathway could thus lead to both an improved understanding of the disease pathogenesis and a validated biomarker to monitor disease progression and response to treatment. Hypusine is an amino acid found exclusively in the eukaryotic translation factor 5A (eIF5A) family [40]. It is important in nonsense-mediated decay [41] but its role in muscle pathology has not yet been established. It is possible that the increase in hypusine is a reflection of ongoing extensive muscle regeneration in *mdx* mice, especially during early disease stages. It is also know that [42] eIF5A induces inflammatory cytokine cascade and the nitric oxide synthase iNOS and inhibition of eIF5A protects mice from developing diabetes. While the precise role of biliverdin and hypusine in vivo in skeletal muscle in *mdx* is unknown, we postulate that their levels in urine likely reflect inflammation and regeneration in dystrophin deficient skeletal muscle.

## Supporting information

## Acknowledgments

Mathula Thangarajh acknowledges the support of the American Academy of Neurology/American Brain Foundation Clinical Research Training Fellowship (2015 – 2017). Yetrib Hathout acknowledges the support of the Muscular Dystrophy Association (MDA353094). We thank Nathaniel Snyder for helpful discussions about metabolite validation.

## References

1 Matthews E, Brassington R, Kuntzer T, Jichi F, Manzur AY. Corticosteroids for the treatment of Duchenne muscular dystrophy. Cochrane Database Syst Rev. 2016; doi:10.1002/14651858.cd003725.pub4

2 Emery AE. Population frequencies of inherited neuromuscular diseases--a world survey. Neuromuscul Disord. 1991;1: 19–29.

3 Hoffman EP, Brown RH, Kunkel LM. Dystrophin: The protein product of the duchenne muscular dystrophy locus. Cell. 1987;51: 919–928.

4 Koenig M, Hoffman EP, Bertelson CJ, Monaco AP, Feener C, Kunkel LM. Complete cloning of the Duchenne muscular dystrophy (DMD) cDNA and preliminary genomic organization of the DMD gene in normal and affected individuals. Cell. 1987;50: 509–517.

5 McDonald CM, Henricson EK, Abresch RT, Duong T, Joyce NC, Hu F, et al. Long-term effects of glucocorticoids on function, quality of life, and survival in patients with Duchenne muscular dystrophy: a prospective cohort study. Lancet. 2018;391: 451–461.

6 Griggs RC, Herr BE, Reha A, Elfring G, Atkinson L, Cwik V, et al. Corticosteroids in Duchenne muscular dystrophy: major variations in practice. Muscle Nerve. 2013;48: 27–31.

7 Hathout Y, Conklin LS, Seol H, Gordish-Dressman, H, Brown KJ, Morgenroth LP, et al. Serum pharmacodynamic biomarkers for chronic corticosteroid treatment of children. Sci Rep. 2016;6: 31727.

8 Boca SM, Nishida M, Harris M, Rao S, Cheema AK, Gill K, et al. Discovery of Metabolic Biomarkers for Duchenne Muscular Dystrophy within a Natural History Study. PLOS ONE. 2016;11: e0153461.

9 Takeshita E, Komaki H, Tachimori H, Miyoshi K, Yamamiya I, Shimizu-Motohashi Y, et al. Urinary prostaglandin metabolites as Duchenne muscular dystrophy progression markers. Brain Dev. 2018;40: 918–925.

10 Rouillon J, Zocevic A, Leger T, Garcia C, Camadro J-M, Udd B, et al. Proteomics profiling of urine reveals specific titin fragments as biomarkers of Duchenne muscular dystrophy. Neuromuscul Disord. 2014;24: 563–573.

11 Rouillon J, Lefebvre T, Denard J, Puy V, Daher R, Ausseil J, et al. High urinary ferritin reflects myoglobin iron evacuation in DMD patients. Neuromuscul Disord. 2018;28: 564–571.

12 Catapano F, Domingos J, Perry M, Ricotti V, Phillips L, Servais L, et al. Downregulation of miRNA-29, -23 and -21 in urine of Duchenne muscular dystrophy patients. Epigenomics. 2018;10: 875–889.

13 Landis SC, Amara SG, Asadullah K, Austin CP, Blumenstein R, Bradley EW, et al. A call for transparent reporting to optimize the predictive value of preclinical research. Nature. 2012;490: 187–191.

14 Willmann R, De Luca A, Benatar M, Grounds M, Dubach J, Raymackers J-M, et al. Enhancing translation: guidelines for standard pre-clinical experiments in mdx mice. Neuromuscul Disord. 2012;22: 43–49.

15 Heier CR, Damsker JM, Yu Q, Dillingham BC, Huynh T, Van der Meulen JH, et al. VBP15, a novel anti-inflammatory and membrane-stabilizer, improves muscular dystrophy without side effects. EMBO Mol Med. 2013;5: 1569–1585.

16 Smith CA, Want EJ, O’Maille G, Abagyan R, Siuzdak G. XCMS: processing mass spectrometry data for metabolite profiling using nonlinear peak alignment, matching, and identification. Anal Chem. 2006;78: 779–787.

17 Gentleman RC, Carey VJ, Bates DM, Bolstad B, Dettling M, Dudoit S, et al. Bioconductor: open software development for computational biology and bioinformatics. Genome Biol. 2004;5: R80.

18 Kuhl C, Tautenhahn R, Böttcher, C, Larson TR, Neumann S. CAMERA: an integrated strategy for compound spectra extraction and annotation of liquid chromatography/mass spectrometry data sets. Anal Chem. 2012;84: 283–289.

19 Bolstad BM, Irizarry RA, Astrand M, Speed TP. A comparison of normalization methods for high density oligonucleotide array data based on variance and bias. Bioinformatics. 2003;19: 185–193.

20 Smith CA, O’Maille G, Want EJ, Qin C, Trauger SA, Brandon TR, et al. METLIN: a metabolite mass spectral database. Ther Drug Monit. 2005;27: 747–751.

21 Wishart DS, Jewison T, Guo AC, Wilson M, Knox C, Liu Y, et al. HMDB 3.0—The Human Metabolome Database in 2013. Nucleic Acids Res. 2012;41: D801–D807.

22 Goloborodko AA, Levitsky LI, Ivanov MV, Gorshkov MV. Pyteomics--a Python framework for exploratory data analysis and rapid software prototyping in proteomics. J Am Soc Mass Spectrom. 2013;24: 301–304.

23 Zhou B, Wang J, Ressom HW. MetaboSearch: tool for mass-based metabolite identification using multiple databases. PLoS One. 2012;7: e40096.

24 Frański R, Gierczyk B, Popenda Ł, Kasperkowiak M, Pędzinski T. Identification of a biliverdin geometric isomer by means of HPLC/ESI-MS and NMR spectroscopy. Differentiation of the isomers by using fragmentation “in-source.” Monatsh Chem. 2018;149: 995–1002.

25 Chen C, Li Z, Huang H, Suzek BE, Wu CH, UniProt Consortium. A fast Peptide Match service for UniProt Knowledgebase. Bioinformatics. 2013;29: 2808–2809.

26 Matsuo M, Shirakawa T, Awano H, Nishio H. Receiver operating curve analyses of urinary titin of healthy 3-y-old children may be a noninvasive screening method for Duchenne muscular dystrophy. Clin Chim Acta. 2018;486: 110–114.

27 Awano H, Matsumoto M, Nagai M, Shirakawa T, Maruyama N, Iijima K, et al. Diagnostic and clinical significance of the titin fragment in urine of Duchenne muscular dystrophy patients. Clin Chim Acta. 2018;476: 111–116.

28 Robertson AS, Majchrzak MJ, Smith CM, Gagnon RC, Devidze N, Banks GB, et al. Dramatic elevation in urinary amino terminal titin fragment excretion quantified by immunoassay in Duchenne muscular dystrophy patients and in dystrophin deficient rodents. Neuromuscul Disord. 2017;27: 635–645.

29 Hathout Y, Marathi RL, Rayavarapu S, Zhang A, Brown KJ, Seol H, et al. Discovery of serum protein biomarkers in the mdx mouse model and cross-species comparison to Duchenne muscular dystrophy patients. Hum Mol Genet. 2014;23: 6458–6469.

30 Antoury L, Hu N, Balaj L, Das S, Georghiou S, Darras B, et al. Analysis of extracellular mRNA in human urine reveals splice variant biomarkers of muscular dystrophies. Nat Commun. 2018;9: 3906.

31 Okinaka S, Kumagai H, Ebashi S, Sugita H, Momoi H, Toyokura Y, et al. Serum Creatine Phosphokinase: Activity in Progressive Muscular Dystrophy and Neuromuscular Diseases. Arch Neurol. American Medical Association; 1961;4: 520–525.

32 Hathout Y, Brody E, Clemens PR, Cripe L, DeLisle RK, Furlong P, et al. Large-scale serum protein biomarker discovery in Duchenne muscular dystrophy. Proc Natl Acad Sci U S A. 2015;112: 7153–7158.

33 Oonk S, Spitali P, Hiller M, Switzar L, Dalebout H, Calissano M, et al. Comparative mass spectrometric and immunoassay-based proteome analysis in serum of Duchenne muscular dystrophy patients. Proteomics Clin Appl. 2016;10: 290–299.

34 Chan MC, Ziegler O, Liu L, Rowe GC, Das S, Otterbein LE, et al. Heme oxygenase and carbon monoxide protect from muscle dystrophy. Skelet Muscle. 2016;6: 41.

35 Sun C, Yang C, Xue R, Li S, Zhang T, Pan L, et al. Sulforaphane alleviates muscular dystrophy in mdx mice by activation of Nrf2. J Appl Physiol. 2015;118: 224–237.

36 Hnia K, Hugon G, Rivier F, Masmoudi A, Mercier J, Mornet, D. Modulation of p38 mitogen-activated protein kinase cascade and metalloproteinase activity in diaphragm muscle in response to free radical scavenger administration in dystrophin-deficient Mdx mice. Am J Pathol. 2007;170: 633–643.

37 Pietraszek-Gremplewicz, K, Kozakowska M, Bronisz-Budzynska, I, Ciesla M, Mucha O, Podkalicka P, et al. Heme Oxygenase-1 Influences Satellite Cells and Progression of Duchenne Muscular Dystrophy in Mice. Antioxid Redox Signal. 2018;29: 128–148.

38 Ryter SW, Morse D, Choi AMK. Carbon monoxide: to boldly go where NO has gone before. Sci STKE. 2004;2004: RE6.

39 Tongers J, Fiedler B, König, D, Kempf T, Klein G, Heineke J, et al. Heme oxygenase-1 inhibition of MAP kinases, calcineurin/NFAT signaling, and hypertrophy in cardiac myocytes. Cardiovasc Res. 2004;63: 545–552.

40 Park MH. The post-translational synthesis of a polyamine-derived amino acid, hypusine, in the eukaryotic translation initiation factor 5A (eIF5A). J Biochem. 2006;139: 161–169.

41 Hoque M, Park JY, Chang Y-J, Luchessi AD, Cambiaghi TD, Shamanna R, et al. Regulation of gene expression by translation factor eIF5A: Hypusine-modified eIF5A enhances nonsense-mediated mRNA decay in human cells. Translation (Austin). 2017;5: e1366294.

42 Maier B, Ogihara T, Trace AP, Tersey SA, Robbins RD, Chakrabarti SK, et al. The unique hypusine modification of eIF5A promotes islet beta cell inflammation and dysfunction in mice. J Clin Invest. 2010;120: 2156–2170.

