## Supplementary material for "High-resolution mass spectrometry-based non-targeted metabolomic discovery of disease and glucocorticoid biomarkers in an animal model of muscular dystrophy"

### Slide 1
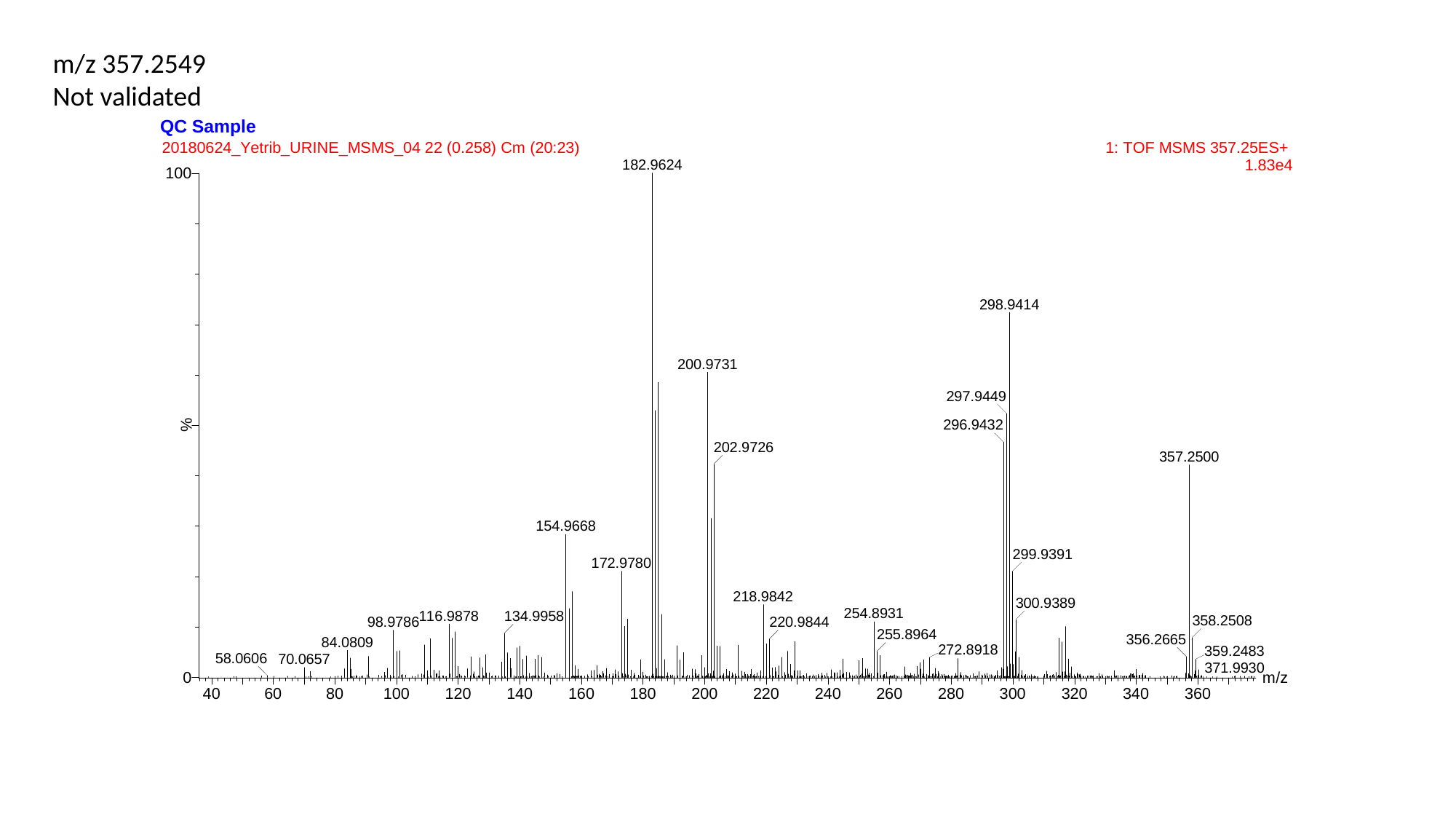

m/z 357.2549
Not validated

### Slide 2
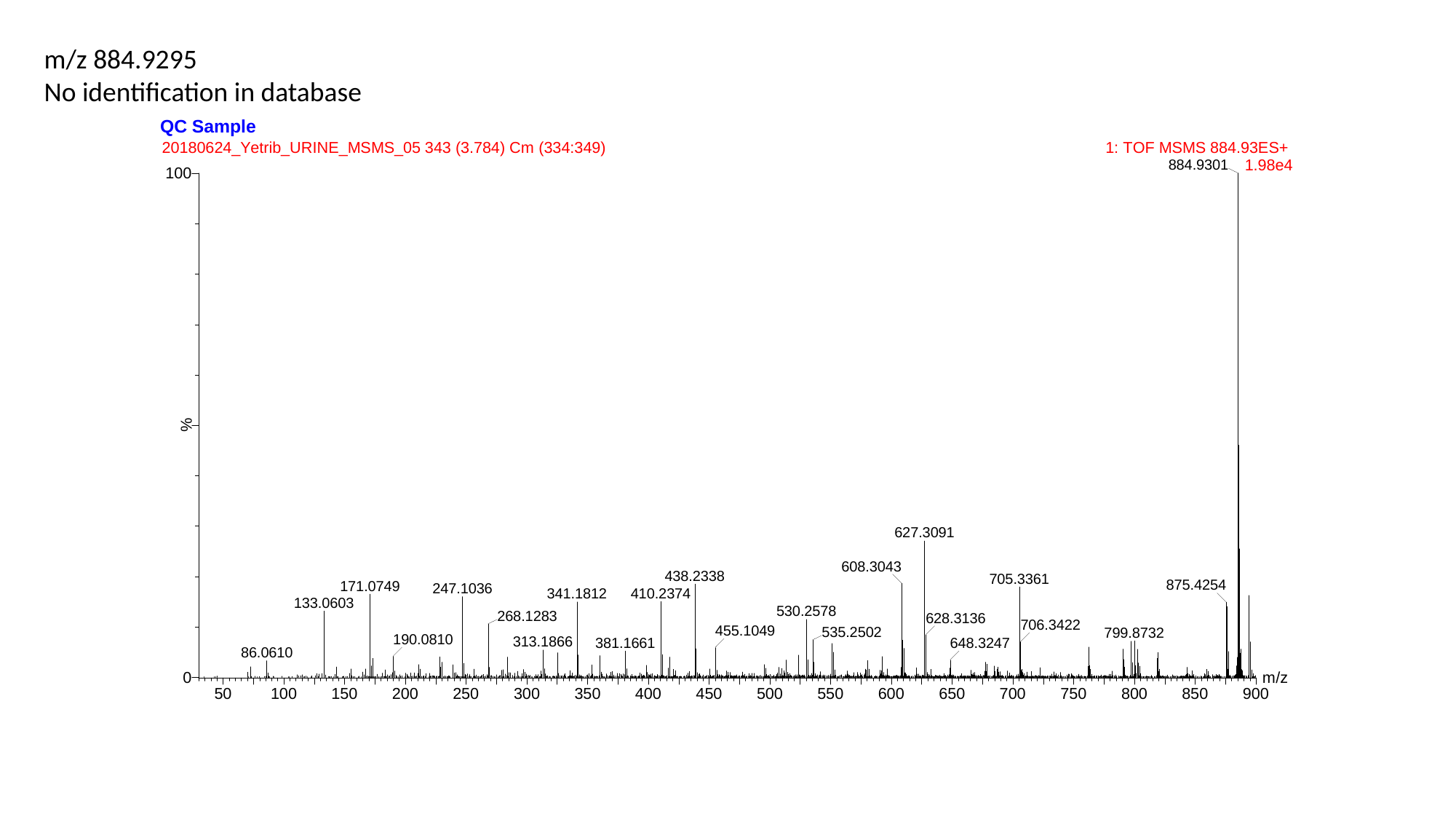

m/z 884.9295
No identification in database

### Slide 3
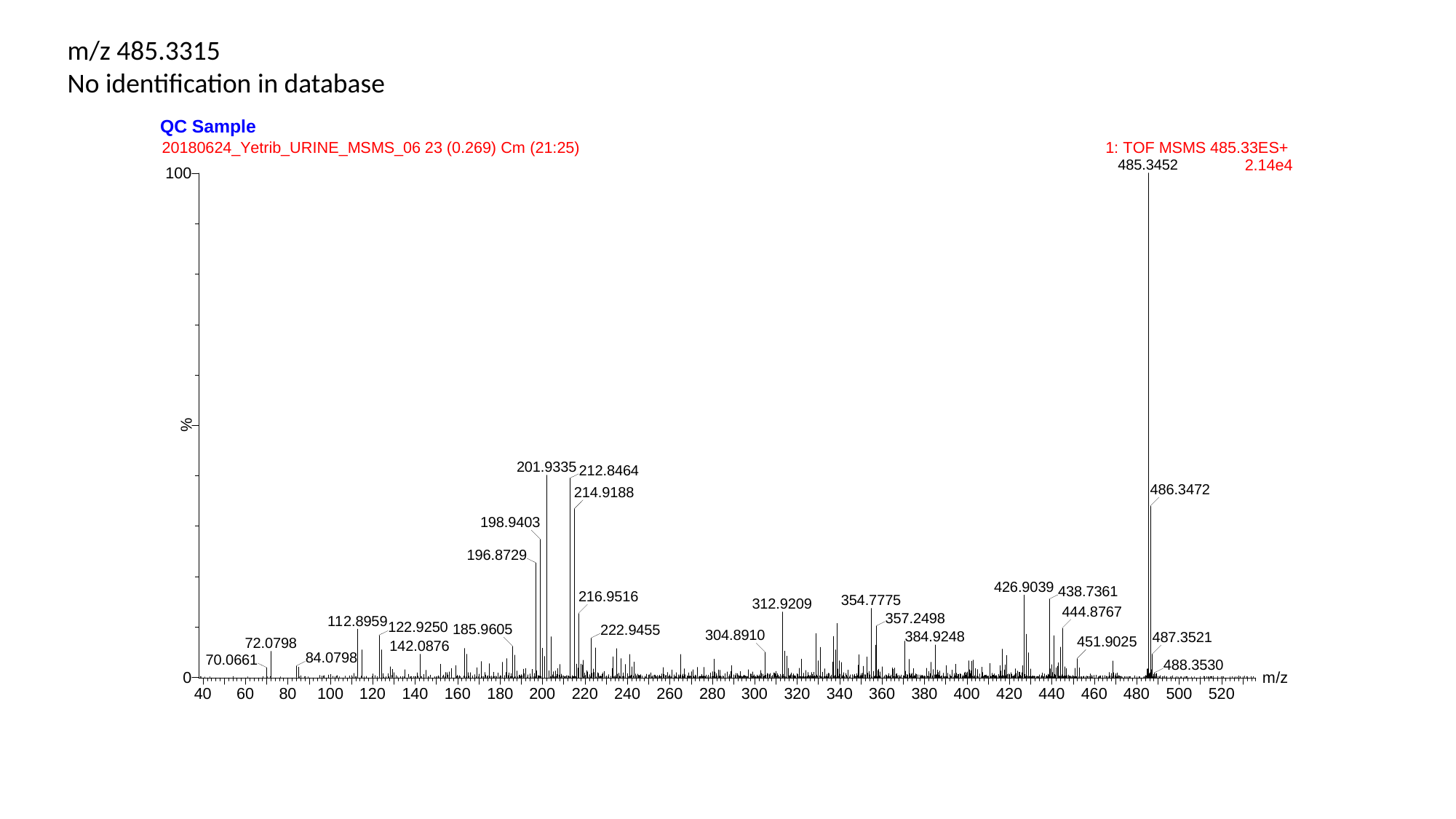

m/z 485.3315
No identification in database

### Slide 4
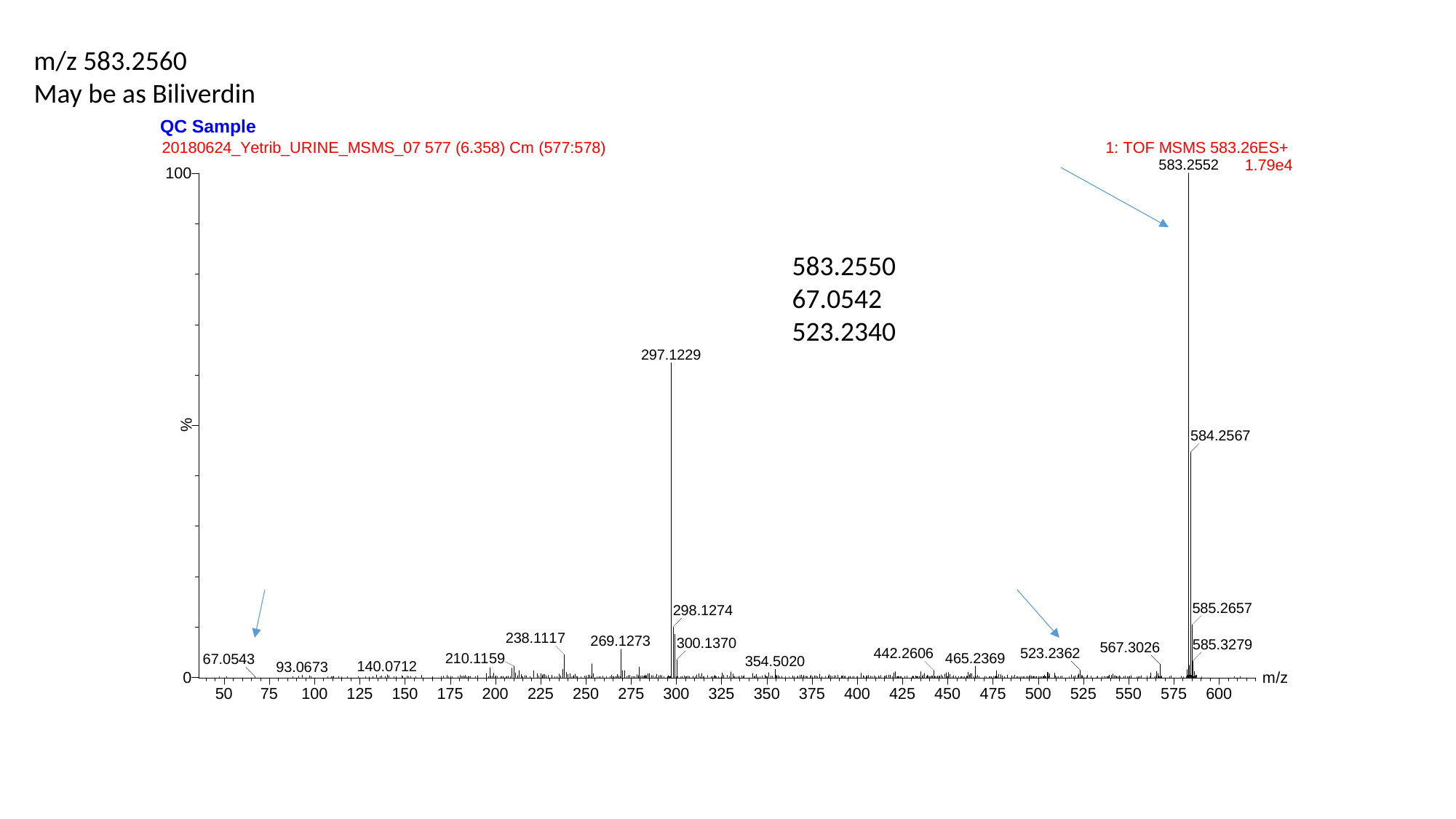

m/z 583.2560
May be as Biliverdin
583.2550
67.0542
523.2340

### Slide 5
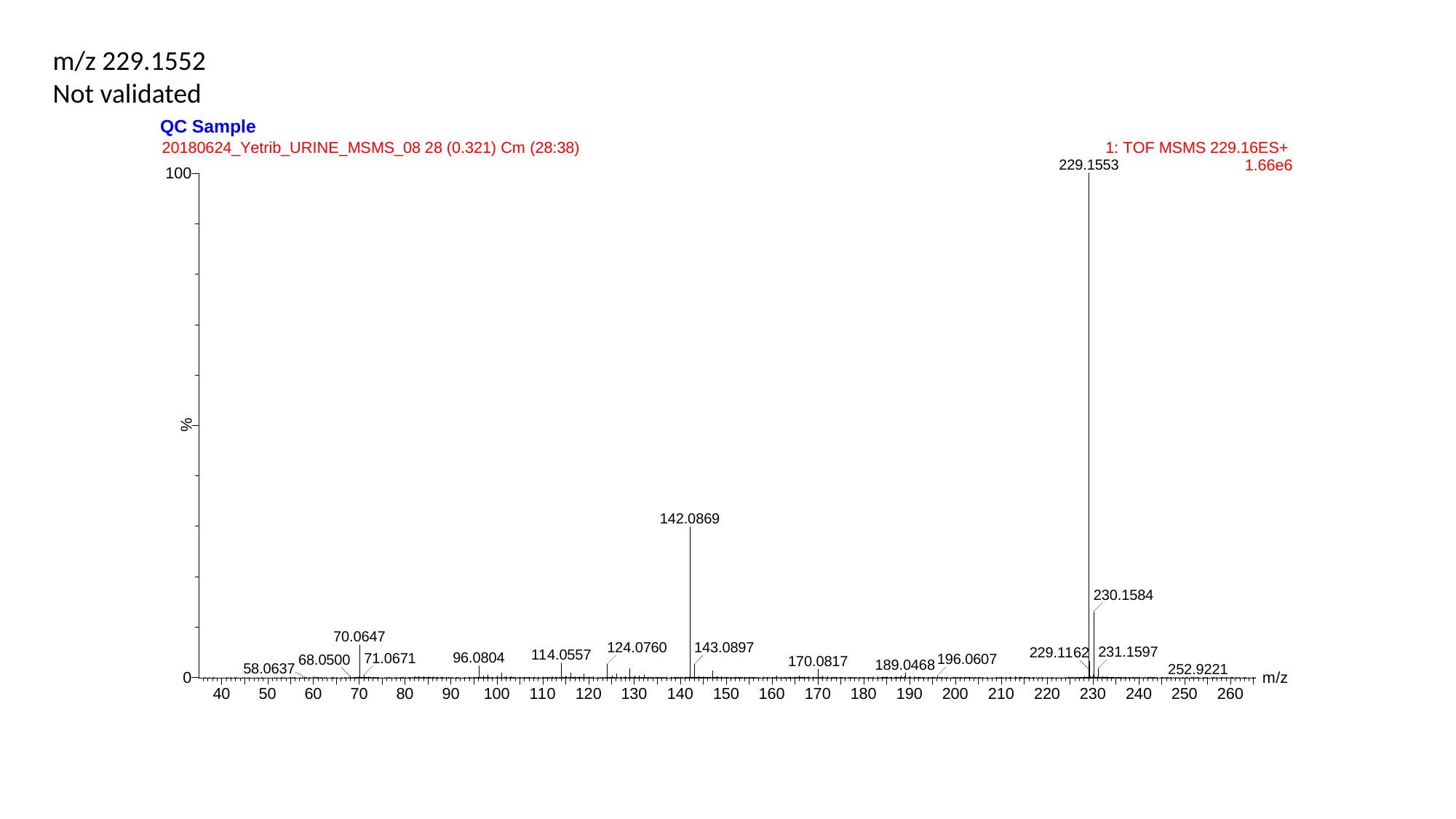

m/z 229.1552
Not validated

### Slide 6
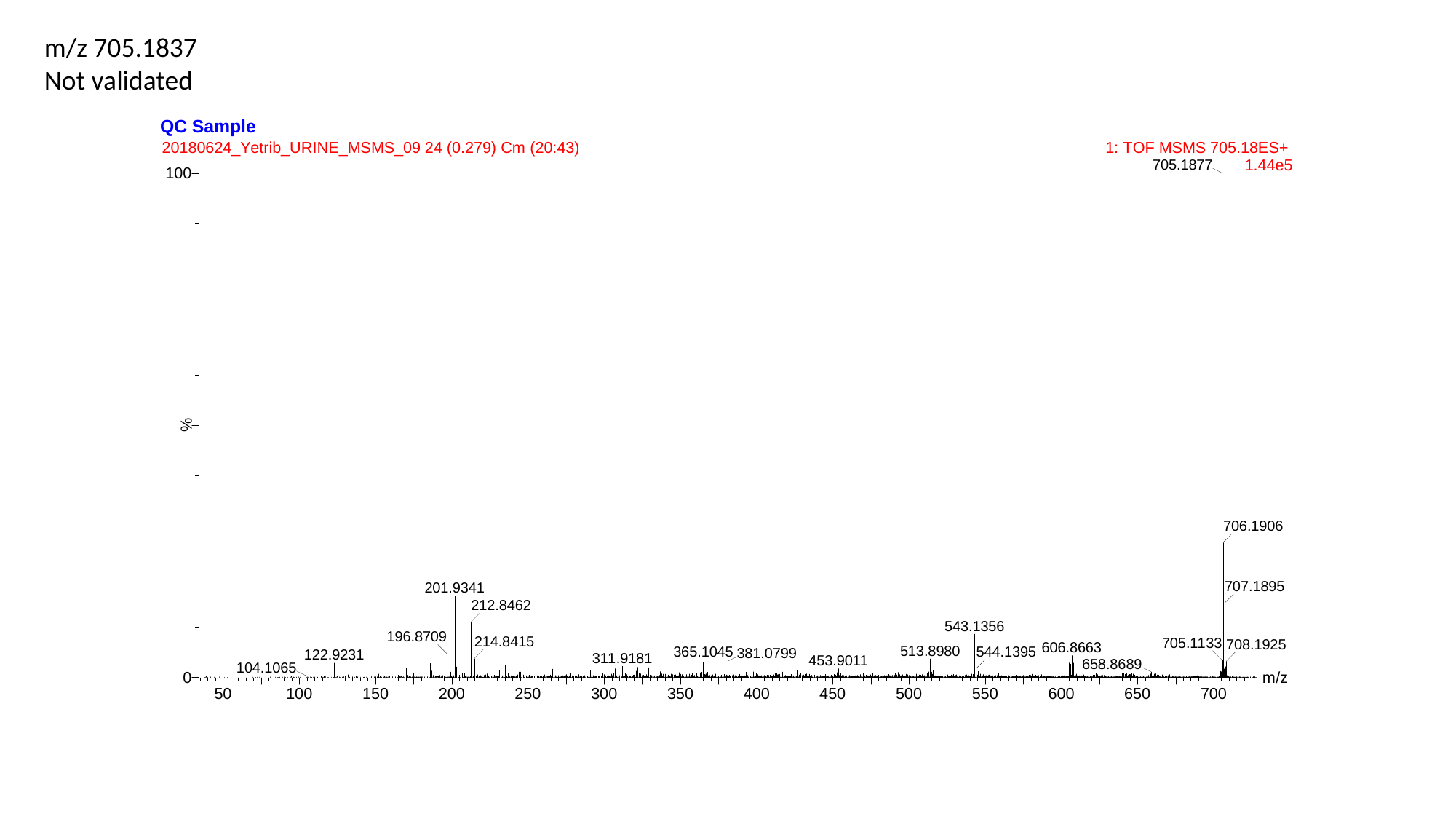

m/z 705.1837
Not validated

### Slide 7
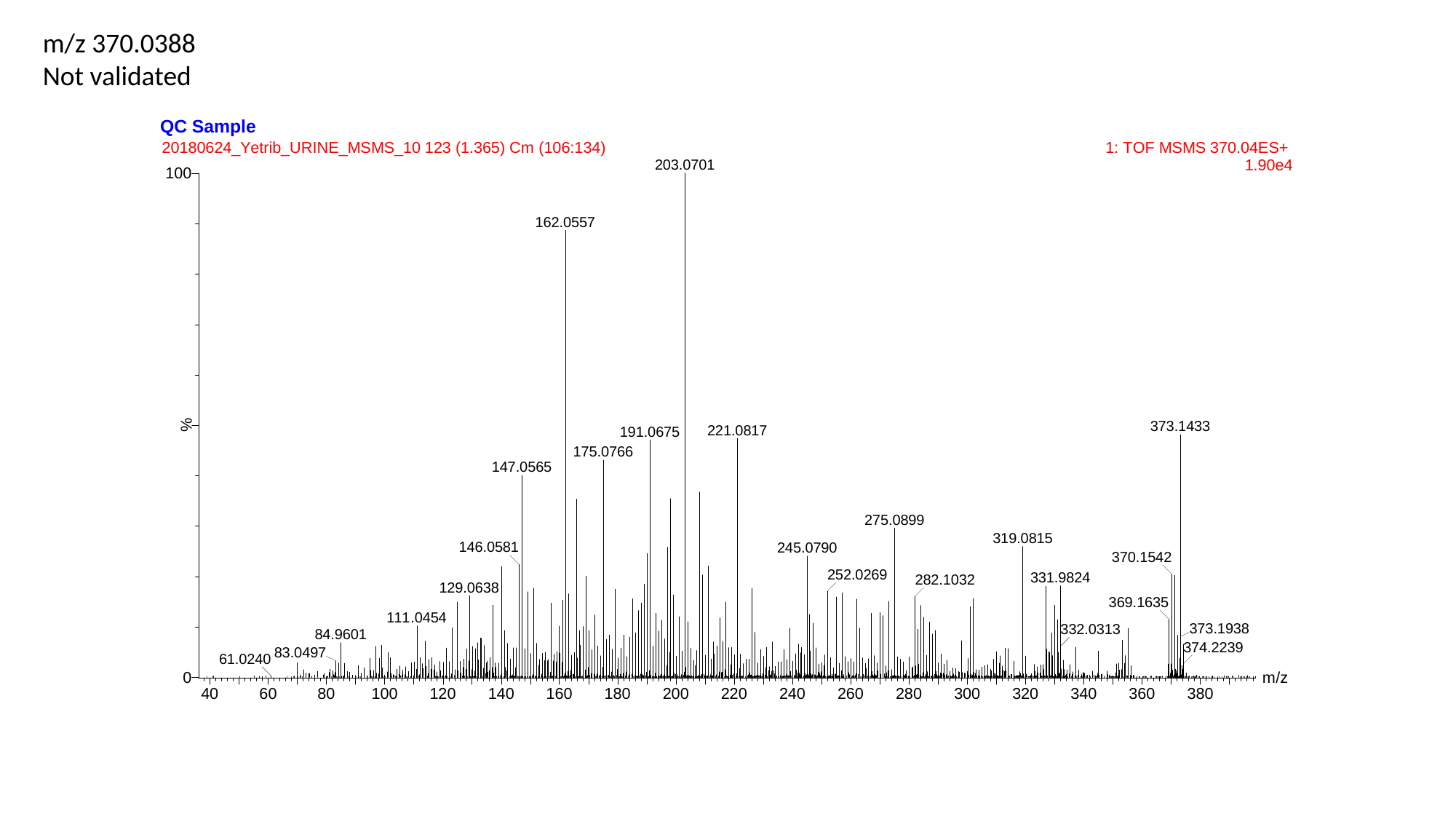

m/z 370.0388
Not validated

### Slide 8
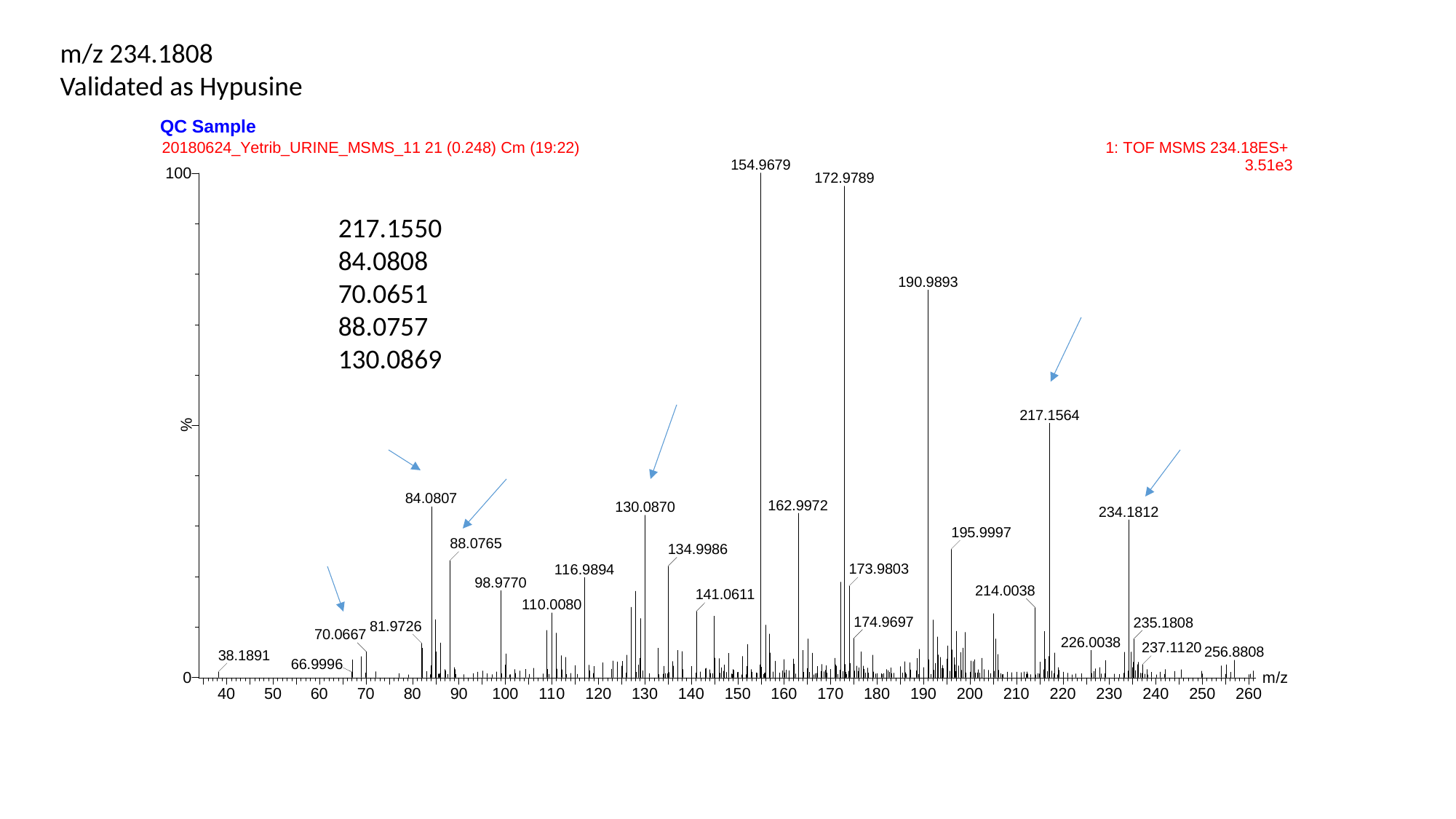

m/z 234.1808
Validated as Hypusine
217.1550
84.0808
70.0651
88.0757
130.0869

### Slide 9
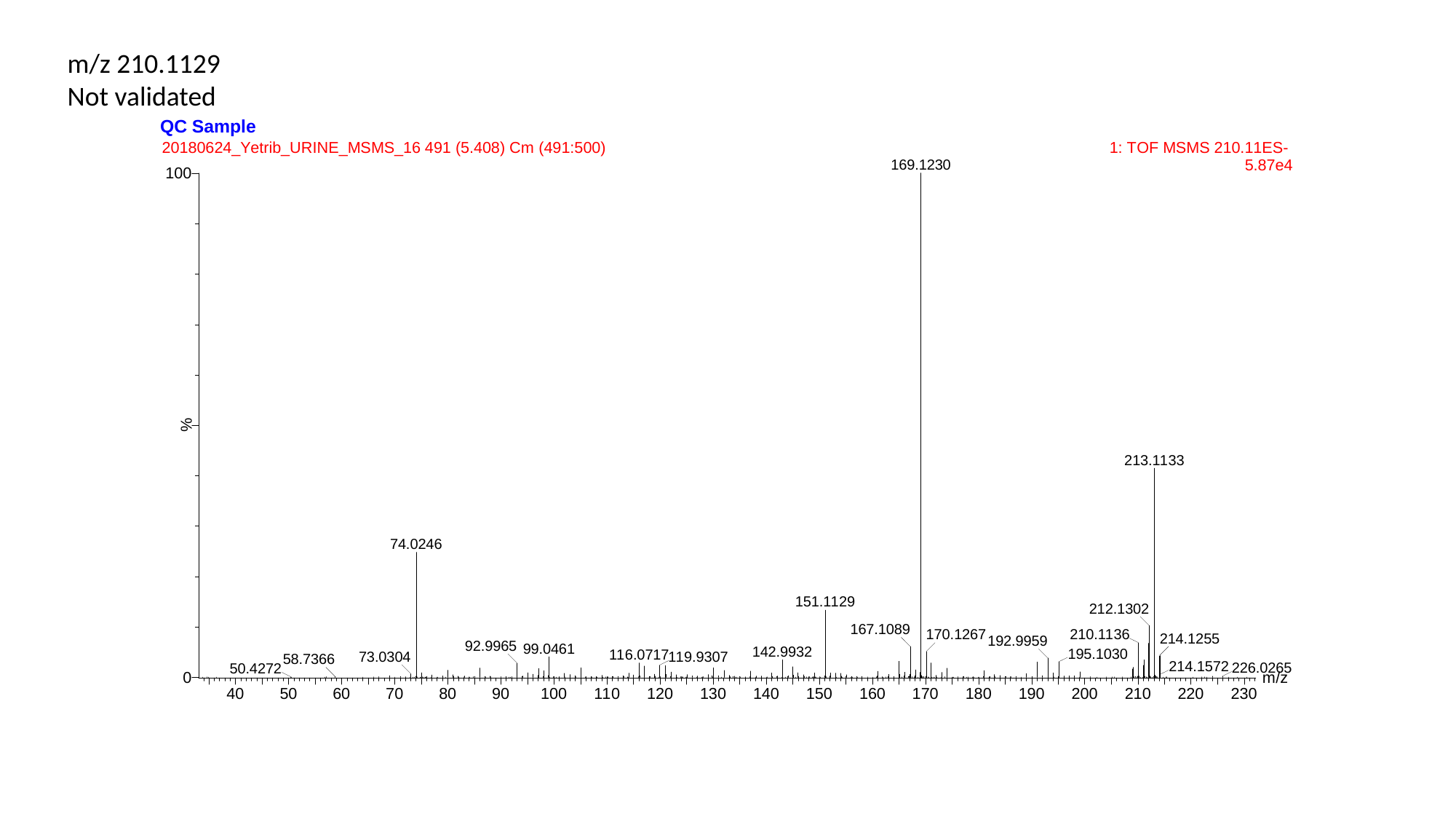

m/z 210.1129
Not validated

### Slide 10
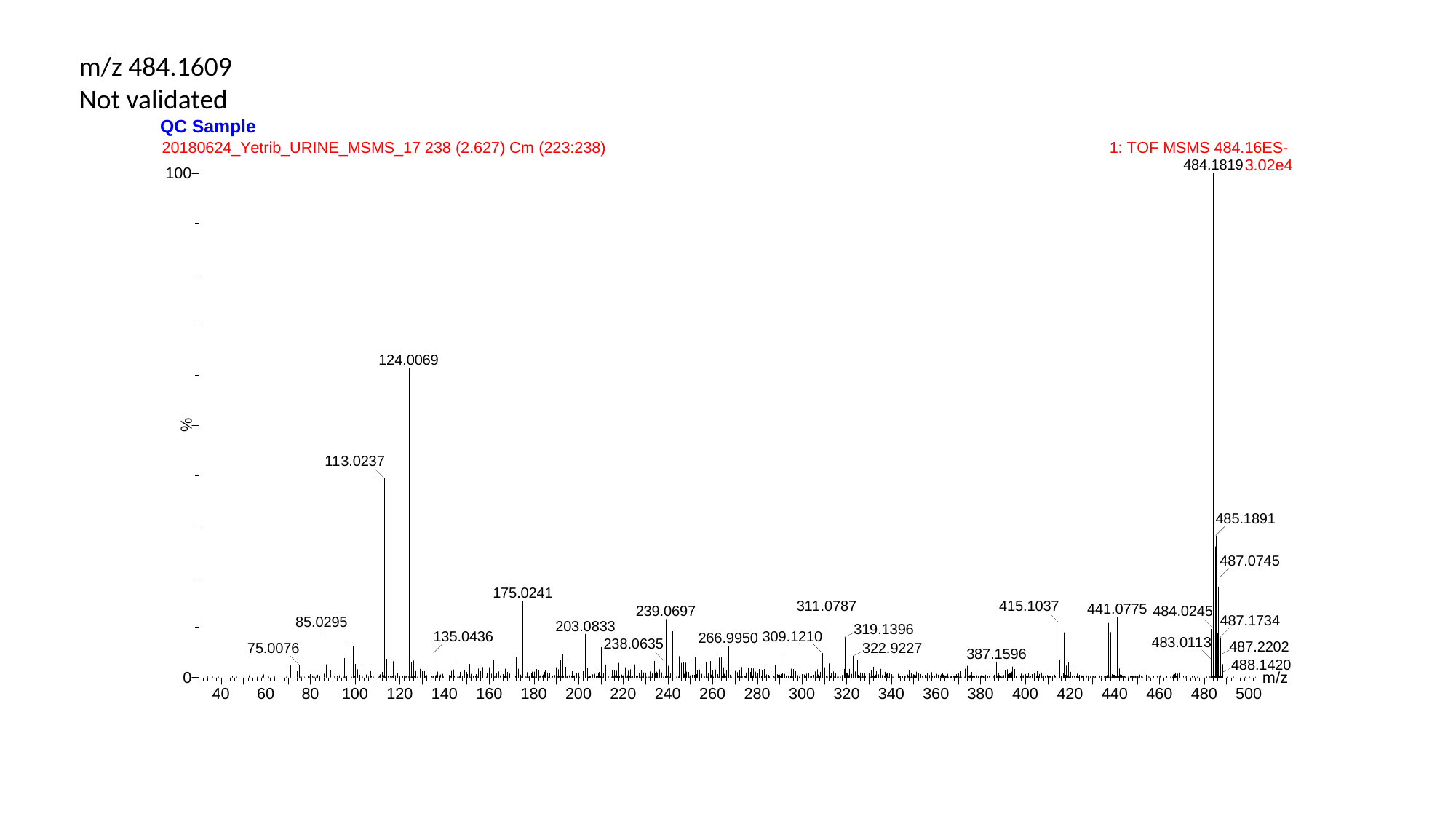

m/z 484.1609
Not validated

### Slide 11
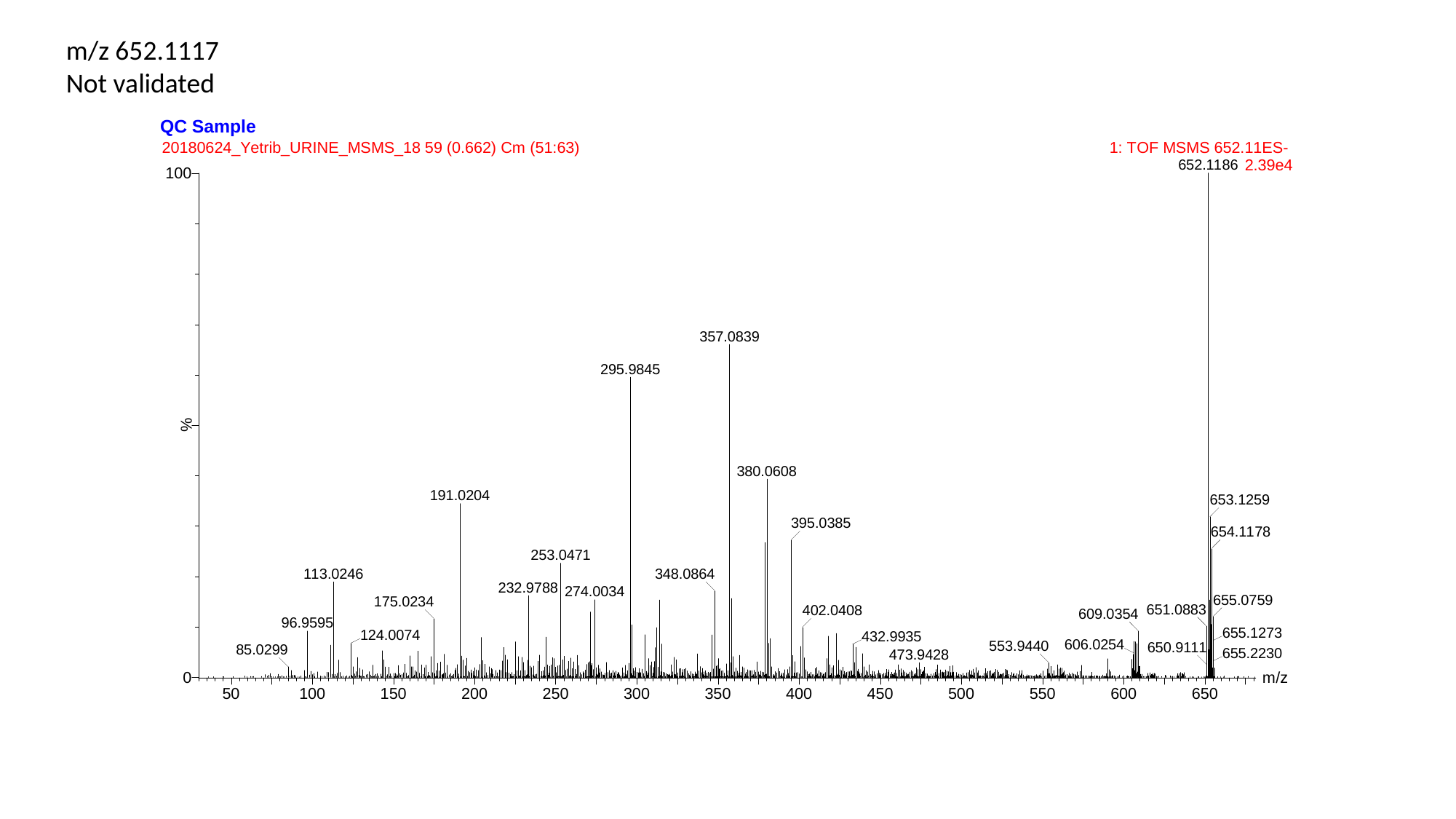

m/z 652.1117
Not validated
