## Supplementary figures and images for "High-resolution mass spectrometry-based non-targeted metabolomic discovery of disease and glucocorticoid biomarkers in an animal model of muscular dystrophy"

### Supplementary file 4

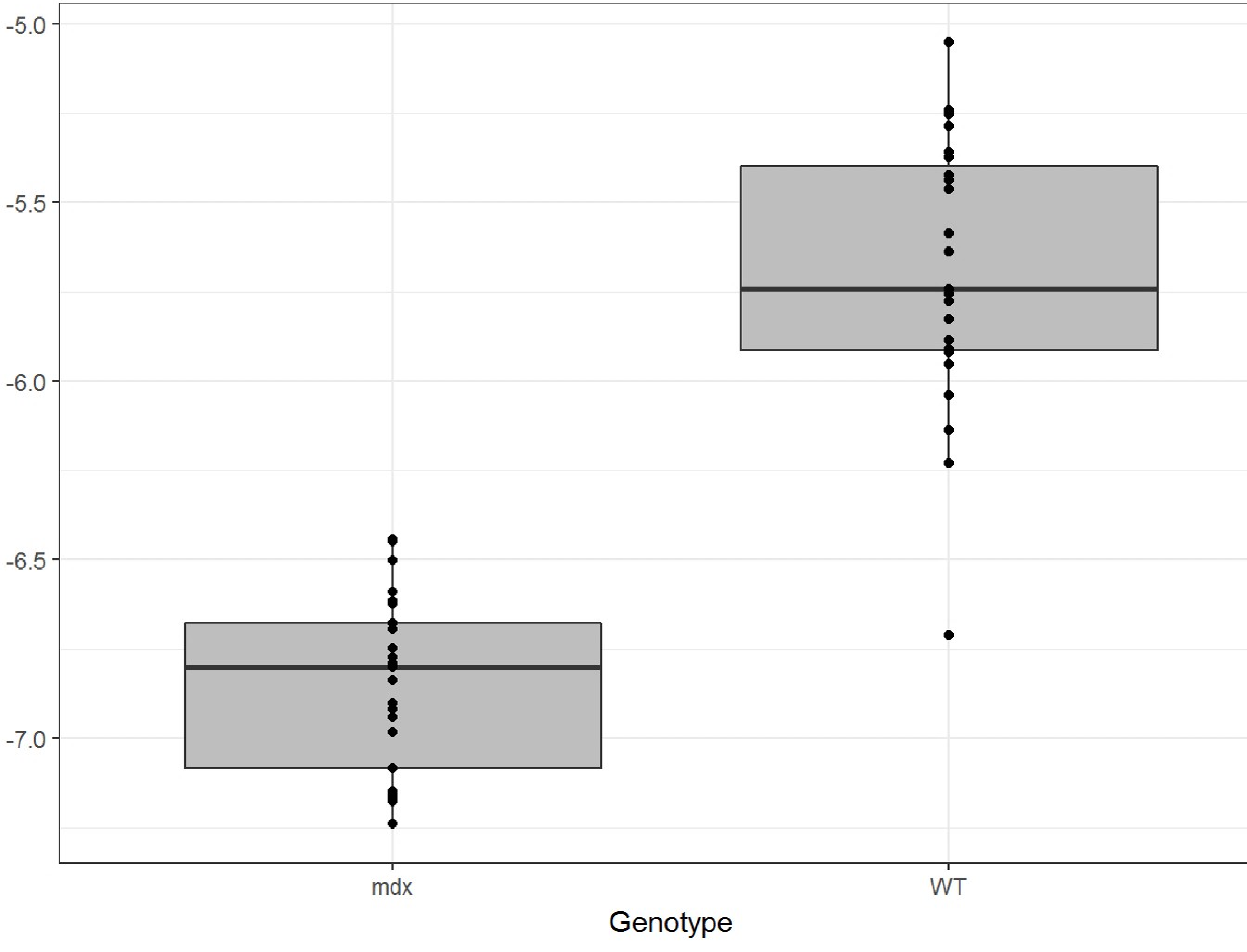

### Supplementary file 5

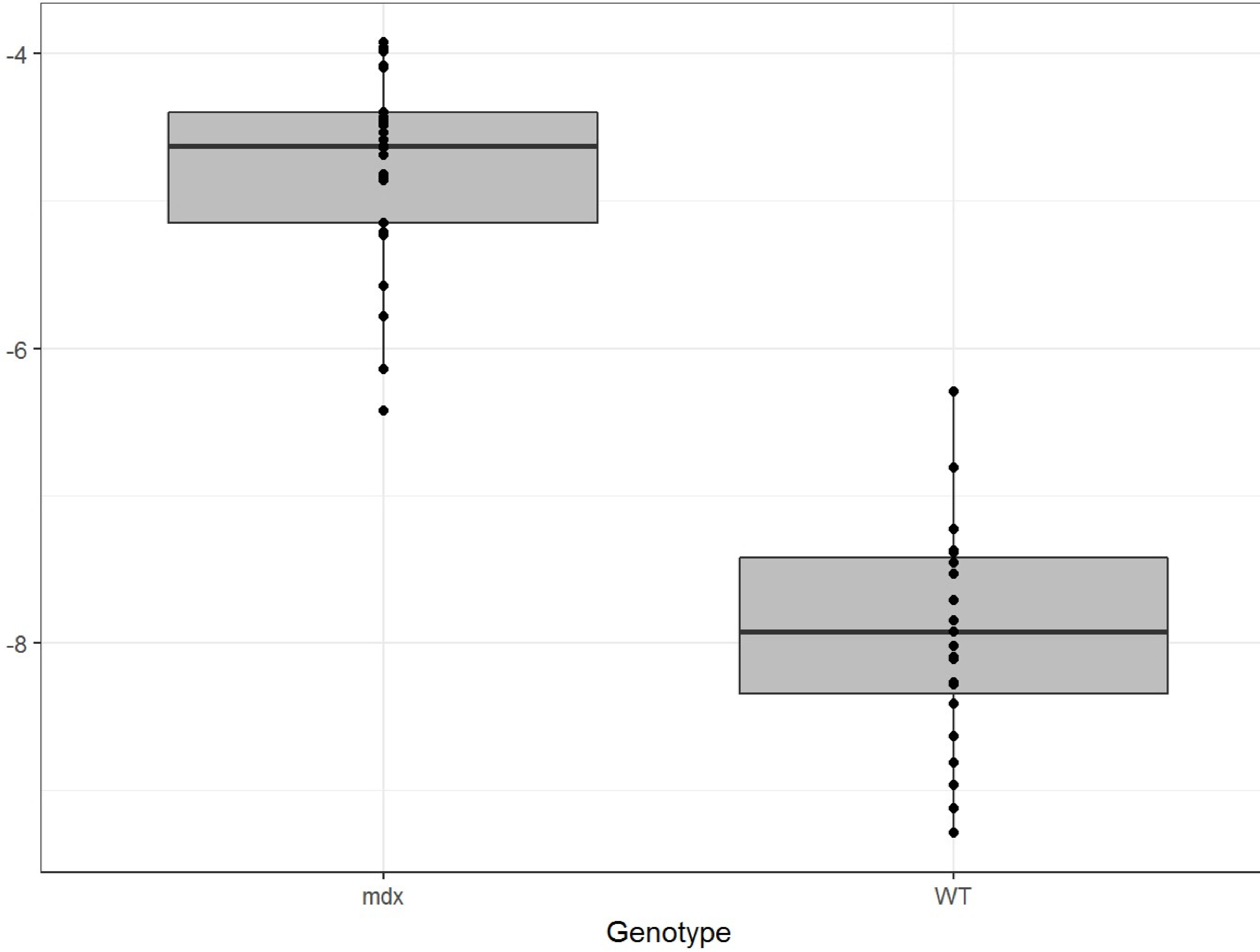
